# Brain connectivity dynamics during social interaction reflect social network structure

**DOI:** 10.1101/096420

**Authors:** Ralf Schmälzle, Matthew Brook O’Donnell, Javier O. Garcia, Christopher N. Cascio, Joseph Bayer, Danielle S. Bassett, Jean Vettel, Emily B. Falk

## Abstract

Social ties are crucial for humans. Disruption of ties through social exclusion has a marked effect on our thoughts and feelings; however, such effects can be tempered by broader social network resources. Here, we use functional magnetic resonance imaging data acquired from 80 male adolescents to investigate how social exclusion modulates functional connectivity within and across brain networks involved in social pain and understanding the mental states of others (i.e., mentalizing). Furthermore, using objectively logged friendship network data, we examine how individual variability in brain reactivity to social exclusion relates to the density of participants’ friendship networks, an important aspect of social network structure. We find increased connectivity within a set of regions previously identified as a mentalizing system during exclusion relative to inclusion. These results are consistent across the regions of interest as well as a whole-brain analysis. Next, examining how social network characteristics are associated with task-based connectivity dynamics, participants who showed greater changes in connectivity within the mentalizing system when socially excluded by peers had less dense friendship networks. This work provides novel insight to understand how distributed brain systems respond to social and emotional challenges, and how such brain dynamics might vary based on broader social network characteristics.

---

Humans are fundamentally motivated to connect with others, spending considerable time and energy investing in social relationships. A lack of social connection due to social isolation has a significant negative impact on health and well-being (1, 2), with risks comparable to those associated with lifelong smoking (3). In fact, the acute disruption of ties through social exclusion has a marked effect on our thoughts and feelings. In contrast, the effects of stress can be tempered by broader social network resources (4–6); as such, brain dynamics may shape the types of networks that individuals inhabit, and different social environments may change the way that individuals approach stressful events (7).

Previous neuroimaging studies have investigated the neural substrates of social exclusion by using experimental designs where participants are socially isolated through rejection by others; one frequently used task is Cyberball, a virtual ball-tossing game where the experimental participant is excluded from the game and receives no ball throws from other players (8, 9). Research on social exclusion has identified the role of two proposed networks: (1) a social pain system, associated with distress during exclusion, (10, 11) characterized by enhanced activations in the anterior cingulate cortex (ACC) and the anterior insula (aINS); and (2) a mentalizing system with consistent activity in the dorsal and ventral medial prefrontal cortex (mPFC), precuneus, and bilateral temporo-parietal junction (TPJ) (12, 13). Although described rhetorically as two cohesive networks, limited research has examined functional interactions, or network dynamics, among the regions during social tasks. A small number of studies have focused on a single hotspot of univariate activation effects in the dACC and assessed its connectivity during social exclusion (14–16); however, no research has examined broader network dynamics during social exclusion. Furthermore, no research has examined how social environments might moderate brain network dynamics in the face of social interaction. This limits our ability to draw conclusions about how brain network and social network dynamics might underlie human processing of social interactions.

To this end, we first capitalize on recent advancements in network neuroscience to study the functional connectivity relationships among multiple brain regions (17, 18) during social exclusion and inclusion. While initially focused on the resting state, these novel techniques have recently been extended to assess the dynamics of connectivity patterns during task performance (19–26). Taking a dynamic network neuroscience perspective, we set out to holistically examine the social pain and mentalizing networks during social exclusion, as well as broader network dynamics across the whole brain as a function of social exclusion and inclusion.

Second, we examine how network connectivity during social exclusion relates to an individual’s social network structure (27, 28). Sociologists have long shown that social network variables can characterize the social structure in which we are embedded and that social network structure can explain important outcomes ranging from measures of happiness to our susceptibility to disease (29–31); likewise, individual differences in personality can shape the structure of social networks (32–35). Building from recent advances in computational social science (36), social network analysis can objectively characterize social network structures related to brain dynamics (27). Specifically, we focus on the density of an individual’s ego-network: a dense network indicates a participant whose friends are also friends with one another, while a sparse network indicates a participant whose friends do not know each other. Perhaps most significantly, dense ego-networks are more close-knit (37, 38) and less likely to include diverse communities. Thus, the friends of the ego are more likely to know, interact, and share with each other, and such close-knit groups may confer social support (39, 40) that may buffer an individual’s response to exclusion. Our analysis focuses on a sample of 80 adolescents males since social ties become especially important during adolescence (41, 42). For each participant, we derived ego-network density using objective logs from the Facebook API. Then, while undergoing fMRI, participants completed Cyberball, a virtual ball-tossing game used to experimentally study the effects of social exclusion (8, 9). An overview of the analysis is shown in Figure 1. We tested the prediction that connectivity during exclusion relative to inclusion would increase in regions of the social pain and mentalizing networks derived from metaanalyses of previous research in Neurosynth (38). We also explored whether connectivity between these two networks would increase during exclusion relative to inclusion. To confirm the effects within these theoretically relevant networks, we also conducted a whole-brain analysis based on a large-scale cortical parcellation comprising 264 regions, including regions that overlap the social pain and mentalizing systems (43). In both analyses, we find that social exclusion is associated with increased connectivity within the regions of the mentalizing network. Furthermore, we link these brain dynamics to individual differences in social network structure and find that participants who show greater connectivity changes in the mentalizing system during social exclusion have less dense friendship networks.

**Figure 1:**
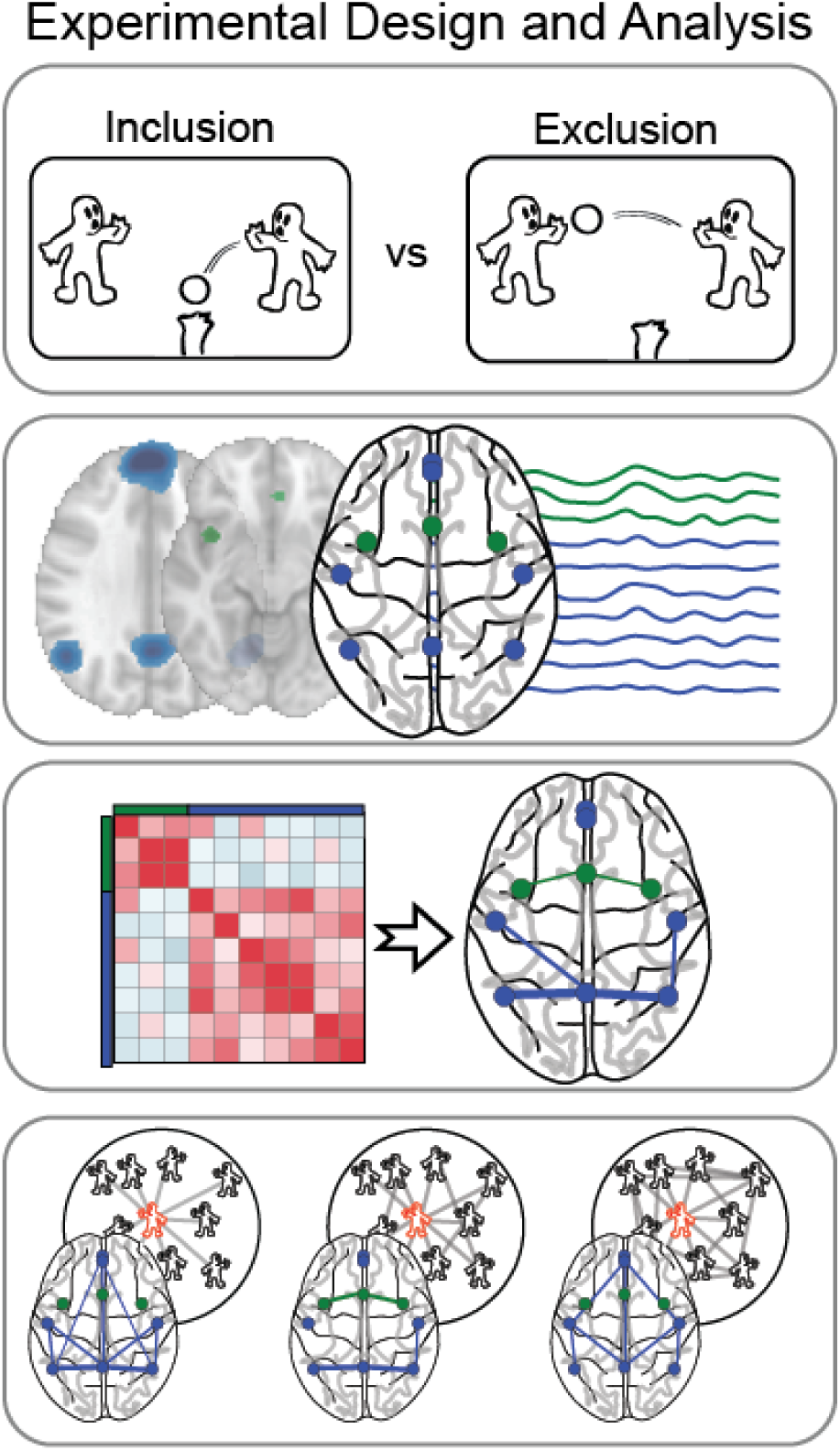
Overview of task and analysis schema. While undergoing fMRI scanning, participants played Cyberball, a virtual game during which they were socially excluded. Using a-priori theorized regions from social pain and mentalizing networks derived from two meta-analyses, we extracted the nodal time series to construct and compare brain network connectivity during social exclusion and inclusion. Finally, we investigated the relationship between each individual’s ego-network density and their brain connectivity during social exclusion.

## Results

### Stronger Connectivity in the Mentalizing Network during Social Exclusion than Inclusion

We first tested the prediction that functional connectivity would increase in social pain and mentalizing systems during exclusion relative to inclusion. We also explored whether connectivity between the two networks changed during exclusion relatively to inclusion. To do so, we examined the functional connectivity between regional time series during social exclusion and social inclusion within *a-priori* networks for social pain processing and mentalizing derived from meta-analyses of each of these constructs (Figure 1; also see Online Methods). We computed connectivity matrices as the Pearson correlation between the time series of every pair of nodes in the two networks. The resulting connectivity matrices for each individual were then group-averaged for display in Figure 2. As can be seen in Figure 2, nodes exhibited high connectivity within both the social pain and mentalizing networks, and only weak connectivity across networks during both inclusion and exclusion; this result supports previous research that has postulated that these regions form two cohesive, segregated networks or modules (44).

**Figure 2:**
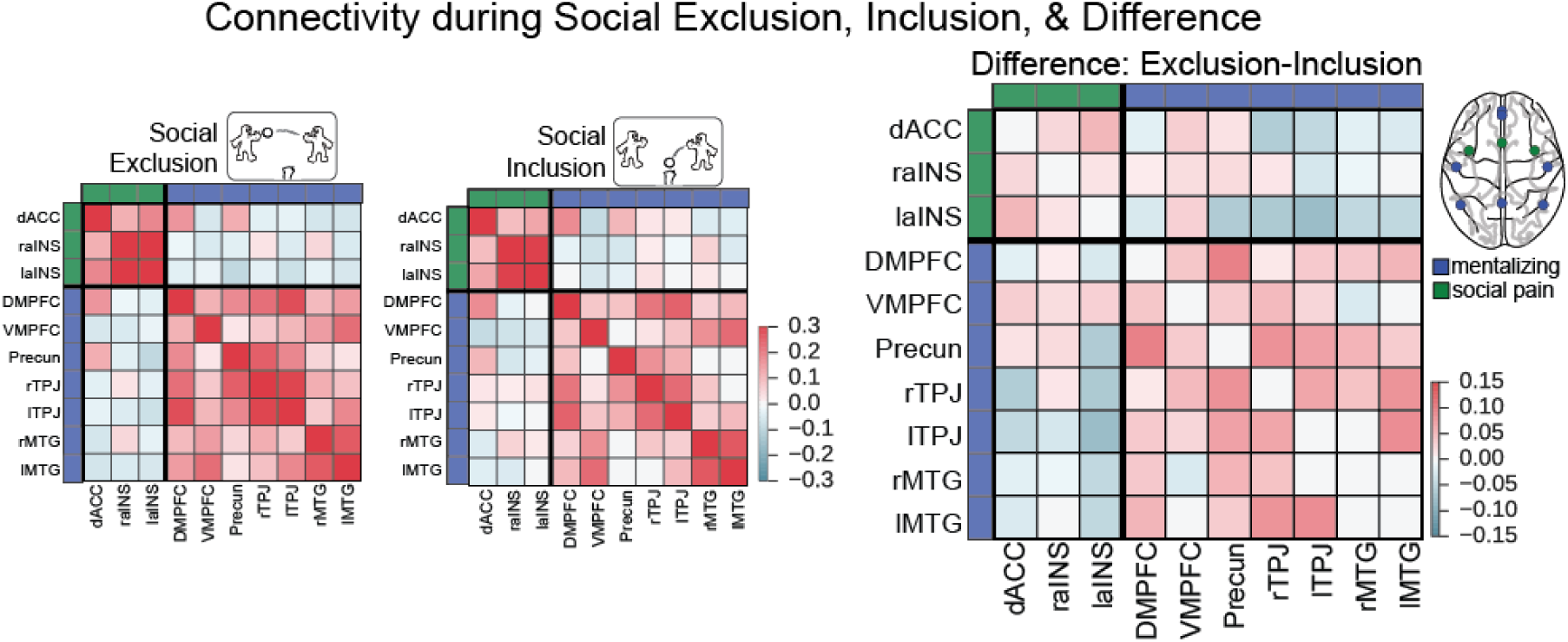
Functional connectivity during social exclusion (A), social inclusion (B), and the difference between exclusion - inclusion (C). For each participant, a functional connectivity matrix was derived as the Pearson correlation coefficient between the time series of every pair of nodes for social exclusion and inclusion, respectively. Group-average functional connectivity matrices were then computed by averaging all participants’ connectivity matrices for each task (after Fisher z-transformation). To isolate the effect of social exclusion, we subtracted the inclusion connectivity matrix from the exclusion connectivity matrix.

Using this functional subdivision of the nodes, we computed the *within-system connectivity* as the mean strength of the functional interactions within each system, and the *between-system connectivity* as the mean of interactions between nodes from different systems (45). We then compared the resulting within- and between-system connectivity values for social inclusion and exclusion with a paired *t*-test. As shown in Figure 3, connectivity significantly increased on average across participants during social exclusion within the mentalizing network, *t(79)* = 3.67, *p* < 0.001, but not within the social pain network or between the social pain and mentalizing networks (*t(79)* = 1.23 and *t(79)* = −1.37, both n.s.). We next asked whether the strength of individual edges, i.e. the connectivity between two nodes, was modulated by exclusion. Similarly, we found that social exclusion was associated with stronger connectivity between edges comprising the mentalizing network (see Supplementary Figure S1). Thus, when people were socially excluded, on average, we observed greater regional connectivity between regions of the mentalizing network, but not between regions of the social pain network or between regions of the two networks.

**Figure 3:**
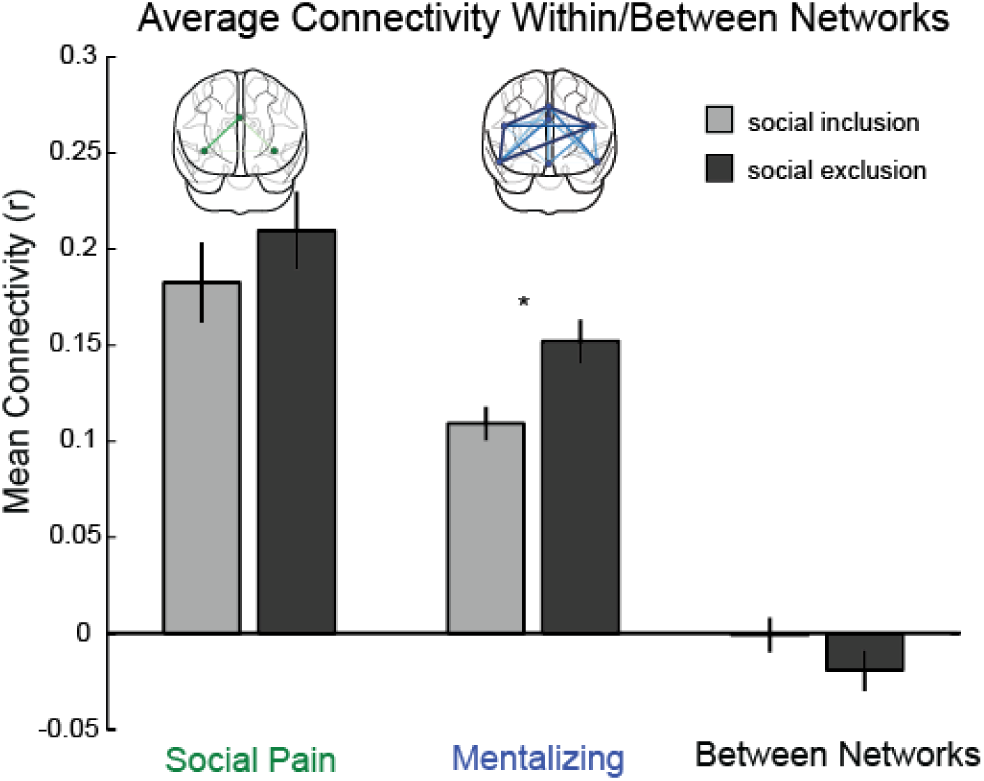
Social exclusion was associated with increased connectivity within the mentalizing network. The strength of recruitment of each network was captured by computing the within-system connectivity as the mean strength of the functional interactions within the social pain (green) and mentalizing network (blue), and the between-system connectivity as the mean of interactions between nodes from the social pain and mentalizing networks (blue nodes to green nodes). Connectivity graphs above each bar illustrate the relative difference in connectivity between social exclusion and inclusion within the respective network. The edge width corresponds to differences in connectivity magnitudes between 0 and 0.1.

To complement our connectivity analysis within *a priori* hypothesized networks, we ran secondary analyses that applied the same analysis pipeline using a whole-brain parcellation to examine both the robustness of our results as well as to identify effects in regions outside of our hypothesized networks of interest. This analysis included 264 brain regions that were previously assigned to 13 large-scale functional brain systems (43), and as before, we computed both within-network connectivity for each of the 13 systems as well as between-network connectivity for all pairs of the networks. Two of these systems have substantial overlap with our *a priori* networks of interest: the default mode network is a more distributed variant of the mentalizing network, while the saliency network is a more distributed variant of the social pain network. This whole-brain analysis also confirmed increased connectivity in the default mode system during social exclusion (*t(79)* = 2.58, *p* = 0.011) as shown in Figure 4. There was no effect within the salience system which overlaps with the social pain network (*t* = −0.86, *p* = 0.39; Figure 4); however, we did observe higher connectivity within the cingulo-opercular system (*t* = 2.08, *p* = 0.041), which overlaps with components of the social pain system. The uncorrected results for all other systems in the Power parcellation were as follows: ventral attention *t(79)* = 0.54, *p* = 0.592; dorsal attention *t(79)* = −1.51, *p* = 0.136; fronto-parietal *t(79)* = 0.99, *p* = 0.325; visual *t(79)* = 2.31, *p* = 0.023; auditory *t(79)* = 2.84, *p* = 0.006; somatosensory/-motor *t(79)* = 1.19; *p* = 0.238; subcortical *t(79)* = −1.47, *p* = 0.146; memory *t(79)* = 1.78, *p* = 0.078; cerebellum *t(79)* = 0.63, *p* = 0.529; uncertain *t(79)* = −1.23, *p* = 0.222.

**Figure 4:**
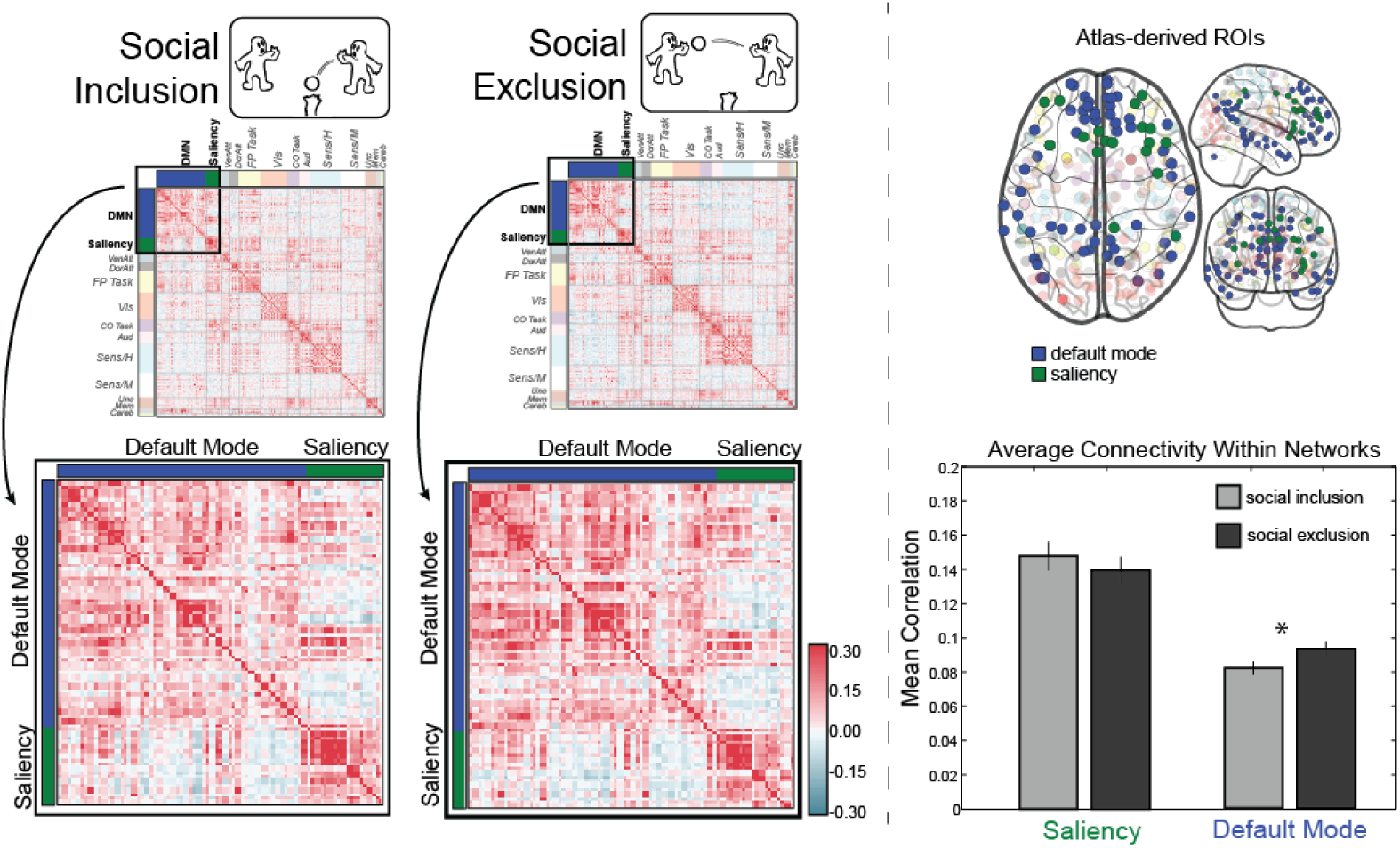
Results from a whole-brain analysis of 264 nodes (Power *et al*, 2011) assigned to 13 systems converged with earlier findings; we observed increased connectivity within the default mode system during social exclusion. For the saliency system, which overlaps with the social pain network, there was no connectivity change during exclusion.

### Individual Differences in Ego-Network Density are linked to Brain Connectivity Effects

Our first set of analyses identified a relationship between connectivity in the mentalizing network and social exclusion across participants in our sample on average, but we hypothesized that the strength of this relationship would be related to individual differences in participants’ social network characteristics. In particular, recent studies suggest a relationship between the mentalizing network and the ‘social landscape’ that people navigate on a daily basis (46, 47); further, psychological factors can shape the structure of social networks (32–35). Consequently, we characterized each individual’s ego-network density from objectively-logged Facebook data, obtained with participants’ consent from the Facebook API. Participants have a denser ego-network when their friends are also friends with one another, versus a sparser network when the participant’s friends are not friends with each other. Especially for adolescent samples, Facebook friendships provide an accurate proxy for offline friendships (48); such social network structures, in turn, can provide a broader window into stable personality differences that are related to important social and emotional outcomes (29).

To determine relationships between the participants’ ego-network data and their brain dynamics during social exclusion and inclusion, we regressed the density of participants’ ego networks onto brain connectivity during social exclusion, controlling for connectivity during social inclusion. Note that we also estimated the correlation coefficient between ego network density and the differences in connectivity during exclusion and inclusion, which yielded the same results. We first entered as connectivity measures the mean strength across nodes from the mentalizing and social pain systems, respectively, but found no significant effects. Next we entered the connectivity of individual edges, i.e., the correlations between the time courses from each pair of nodes. After correcting for multiple comparisons using the false discovery rate (FDR, *q* = 0.05), we found that connectivity between left and right TPJ, two key nodes of the mentalizing network, was negatively related to ego-network density. In other words, individuals who exhibit a stronger TPJ-coupling during exclusion tend to have a less dense friendship network. We did not observe effects between any pairs of nodes from the social pain network, either corrected or uncorrected. Together, these results suggest that brain dynamics during key social experiences such as exclusion may contribute to the type of network structure people occupy. Likewise, the amount of interconnections among friends of the excluded participant’s social network may also influence the impact of exclusion on brain connectivity.

**Figure 5:**
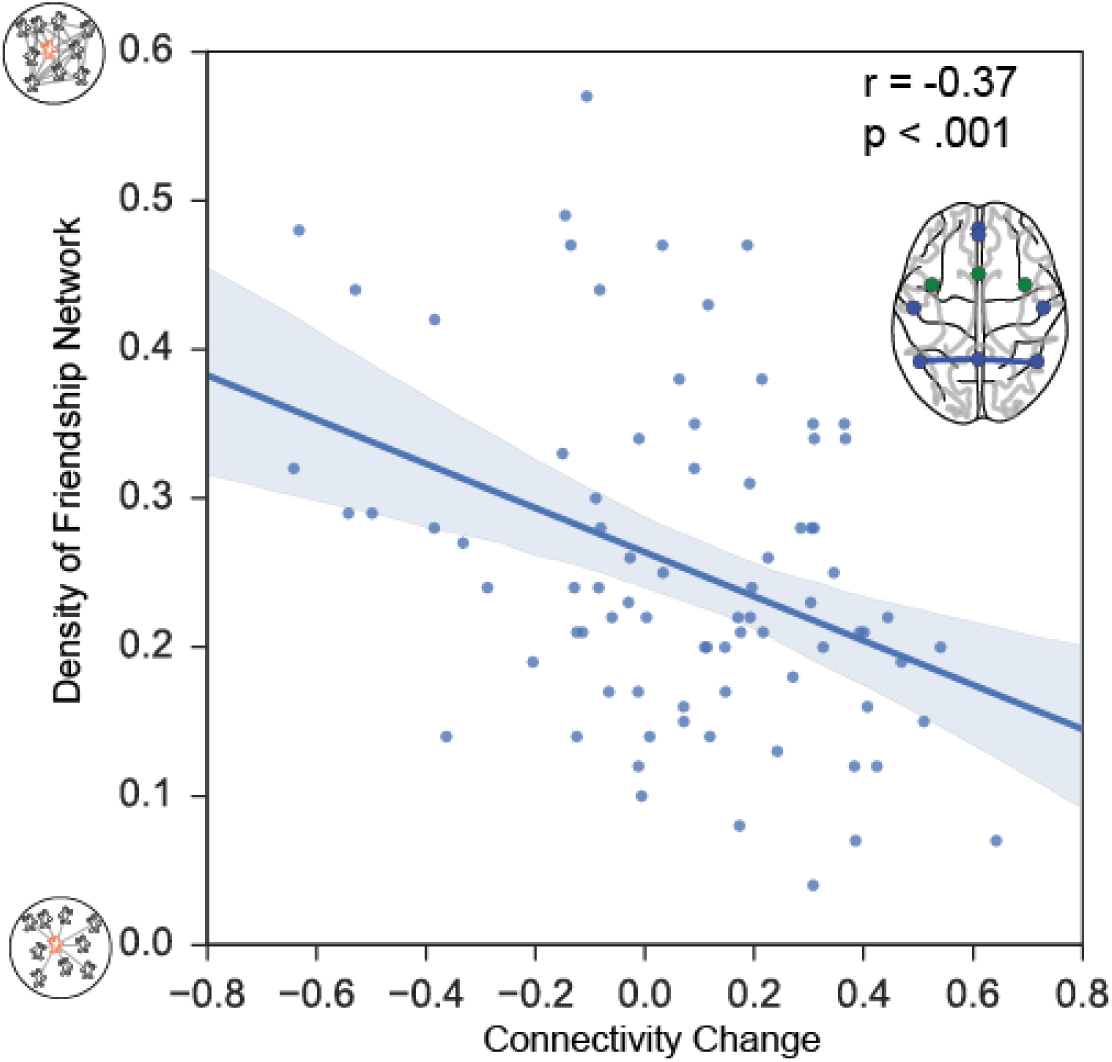
Scatterplot illustrating the significant relationship between the functional connectivity between left and right temporo-parietal junction (TPJ) and the density of a participant’s egofriendship-network. Participants who show stronger regional coupling during exclusion have sparsely connected networks where fewer of their friends are also friends with each other.

## Discussion

Social relationships are a driving force for human behavior and previous research on social exclusion has identified robust involvement of two sets of brain regions activated during social pain and mentalizing. We bring a novel network neuroscience perspective to examine how social experiences may engage and reconfigure these brain networks in support of social cognition, and how individual differences in how people use their brains may correspondingly influence or be influenced by the structure of their social networks.

Specifically, we first tested the prediction that functional connectivity would increase within social pain and mentalizing networks during social exclusion relative to inclusion. We also explored the relationships between the networks during both exclusion and inclusion. Here, we find support for tight coupling of the two proposed networks (social pain and mentalizing) during social inclusion and exclusion. Next, we find that connectivity within the mentalizing system increases during exclusion relative to inclusion. This result is confirmed using a whole brain analysis that identified significant effects within the so-called default mode network. This *atlas-derived* default mode network overlaps with an *a priori* hypothesized mentalizing network.

Previous research has highlighted the role of the social pain system in responding to exclusion (11, 49). Building on this starting point, the few studies examining brain connectivity during social exclusion (14–16, 50) have focused on dACC seed regions as a hotspot of univariate activation effects in the social pain network. Our findings highlight a broader network perspective. Although connectivity is strong within both social pain and mentalizing networks during both social exclusion and inclusion, connectivity increases within the mentalizing system, but not the social pain system, during exclusion relative to inclusion. This first finding highlights the critical value of coordination between regions of the same subnetwork, revealing that withinnetwork instead of between-network interactions were strongly correlated with the social exclusion condition. The second finding complements existing univariate accounts of increased activity within specific nodes of the social pain and mentalizing networks by highlighting a distinction in the temporal dynamics of social pain and mentalizing processes that was not captured by average univariate activation results. Overall, these findings are in line with the wider literature arguing that connectivity and activity provide complementary information (51, 52) and thus can provide complementary insights about cognitive processes.

Functionally, this stronger coupling among mentalizing regions may support considering the intentions of the individuals who are excluding the participant in the social event, or internal ruminations about the relevance of potential ongoing exclusion for one’s broader social relations. In fact, verbal reports from participants who have undergone this type of exclusion show that such thoughts are frequent (8), though not uniformly distributed across people. Indeed, whereas relatively sustained feelings of emotional distress or social pain would logically lead to the type of robust findings previously reported with respect to average activity within the social pain system (10, 11), the connectivity methods used here are also sensitive to thought processes that wax and wane multiple times during the exclusion period, such as fluctuations in mentalizing. Thus, we suggest that increased connectivity in the mentalizing network may tap into such dynamic aspects of the response to social exclusion over a period of time, whereas activity within the social pain system may be relatively sustained over time. One possible function of such a dynamic process may be to make sense of the situation and to reflect more broadly on the meaning of that experience, thereby supporting coping-related functions (53), in contrast to social pain, which may be more sustained throughout the exclusion period.

The robustness of the dynamic fluctuations within the mentalizing network was investigated in a control analysis that compared time courses from the first and second half of each block, confirming our main findings that showed increased within-network connectivity was sustained throughout the first and second halves of each block (first half: *t*(79) = 2.23; *p* = 0.029, and second half: *t*(79) = 2.57, *p* = 0.012, see Supplementary Figure S2). This suggests that participants were not increasingly mind wandering or otherwise engaging default mode activity over time, but rather that our analysis likely tracks social cognitive processes relevant to the task.

Our social-cognitive interpretation of changing connectivity within the mentalizing system as a means of coping with exclusion is also supported by the fact that this effect covaries with participants’ broader social network structures. Specifically, participants who showed increased connectivity during exclusion relative to inclusion between left and right TPJ, two regions of the mentalizing system, also had less dense social network structures. One possibility is that outside of the safety of dense ego-networks, which tend to be more close-knit (37, 38), individuals use greater mentalizing resources, especially during stressful social interactions such as exclusion. This effect is consistent with neurocognitive models that link intrapersonal effects of social exclusion to the interpersonal contexts surrounding individuals during daily life (55). Social exclusion, by its very nature, does not occur in a vacuum. Rather, pre-existing characteristics of one’s social network provide the background or contextual standard from which any new episode of exclusion is understood and compared (54). Within this framework, our measure of social network density can be understood as a social inclination. When excluded, people who interact more with unconnected others may use mentalizing resources differently than those with denser ego-friendship networks. That said, it is also likely that those who use mentalizing resources differently may position themselves differently in their social networks, and that the ways that individuals use their brains during social experiences shapes their preferences and tendencies to occupy different types of network positions. For example, the extant literature suggests that the lack of diverse communities in denser networks may relate to the degree of mentalizing that individuals engage in during social tasks. Indeed, prior work has shown that network brokerage, which is inversely related to density and associated with bridging diverse communities, can moderate mentalizing activity during social decision making (55). The present findings substantially extend prior research linking personality to the shape of social networks (32–35), by showing that individual differences in brain dynamics may underpin such links. In other words, the relationship between brain network dynamics and social network dynamics is likely bidirectional, and the current findings open new avenues to understand both.

Our results add to a growing literature that links individual differences and social network properties, but they also bring novel neuroscientific methods to bear in characterizing individual differences. For example, past research has linked social personality tendencies, such as extraversion, self-monitoring, and cognitive empathy (related to mentalizing) to personal network structure (34, 56). Our results also complement work across other social species, including research with macaques (57). Researchers have studied the structure and connectivity of brain regions that belong to the DMN, including the posterior superior temporal sulcus (pSTS), which parallels our TPJ result. These regions are modulated on the basis of social network characteristics, such as network size and position in a social hierarchy (58). Our data complement these findings by showing that properties of an individual’s social networks are associated with intrapersonal reactions to exclusion, which then might reciprocally influence the interpersonal relationships that underlie these social networks. Taken together, these results also highlight the need to jointly consider structural characteristics of personal networks in combination with cognitive responses to acute episodes of exclusion.

In sum, we find that social exclusion is associated with increased connectivity within the mentalizing system. Furthermore, we explore whether the impact of social exclusion on brain connectivity relates to the structure of participants’ friendship networks, finding that participants with sparser friendship networks show increased connectivity within key brain systems when excluded. These findings demonstrate that networked brain dynamics during social-cognitive tasks can provide relevant information in addition to univariate activity. In particular, our connectivity analysis highlights the fundamental importance of the mentalizing system in responding to socially salient events, and it indicates how a participant’s social network structure may capture critical differences between individuals in how social exclusion influences these key dynamics of the underlying brain networks for social cognition. Together, our connectivity analysis offers novel insights into the neural and social responses to common social experiences such as exclusion and delineates how distributed brain networks respond to powerful socioaffective challenges that manifest in social networks indexed by online media.

## Methods

### Participants

80 neurotypical adolescent males aged 16–17 years were recruited through the Michigan state driver registry database as part of a larger study on peer influences on adolescent driving. Participants met standard MRI safety criteria. In accordance with Institutional Review Board approval from the University of Michigan, legal guardians provided written informed consent, and adolescents provided written assent.

### Social exclusion task

Participants completed the Cyberball game while undergoing functional MRI scanning, which has been validated in a number of behavioral and neuroimaging studies as a reliable way of simulating the experience of social exclusion (10, 59). The Cyberball game consisted of two three-minute rounds and the order of rounds was held constant to preserve the psychological experience across participants. In the inclusion round, the participant and two virtual players received the ball equally often, whereas during the exclusion game the participant and virtual players started out playing the ball, but the participant was then left out after a few throws, simulating social exclusion. After the scan, participants completed a set of questionnaires.

### Social network assessment

In addition to the fMRI tasks, participants also provided information about their social networks. This information was assessed from logged online friendships using the Facebook API (collected in 2011–2013). Following this data acquisition, density was computed for participants’ egocentric networks. In particular, we constructed a friendship network for a given individual (‘ego’), then removed the ego from the network (since all friends are logically connected to it), and computed the density among the remaining nodes (*n*) of friends, assessing the proportion of existing connections (*m*) between friends vs. all possible connections (i.e. *n*\**(n-1)/2* if all of your friends are also friends with each other). In other words, density of the ego-network measures the extent to which participants’ Facebook friends are interconnected. After first removing the ego from each individual’s friendship network, density was computed using NetworkX with the formula:

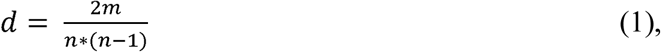

where *m* denotes the number of edges between persons and *n* denotes the number of nodes.

### FMRI acquisition and analysis

Functional images were recorded using a reverse spiral sequence (repetition time = 2000 msec, echo time = 30 msec, flip angle = 90°, 43 axial slices, field of view = 220 mm, slice thickness = 3 mm, voxel size = 3.44 × 3.44 × 3.0 mm). We also acquired in-plane T1-weighted images (43 slices, slice thickness =3 mm, voxel size = 0.86 × 0. 86 × 3.0 mm) and high-resolution T1-weighted images (SPGR, 124 slices, slice thickness = 1.02 × 1.02 × 1.2 mm) for use in coregistration and normalization. Functional data were preprocessed and analyzed using Statistical Parametric Mapping (SPM8, Wellcome Department of Cognitive Neurology, Institute of Neurology, London, UK). The first four volumes were discarded before analysis. Functional images were despiked using the 3dDespike program as implemented in the AFNI toolbox, corrected for slice time acquisition differences, and spatially realigned to the first functional image. Functional and structural images were coregistered using a two-stage procedure. First, in-plane T1 images were registered to the mean functional image. Next, high-resolution T1 images were registered to the in-plane image. Structural images were then skull-stripped and normalized to the skull-stripped MNI template provided by FSL.

### Meta-analytical definition of regions

Nodes for social pain and mentalizing networks were derived based on previous literature: We used Neurosynth (38) to perform two automated metaanalyses of the functional neuroimaging literature on “mentalizing” and “social pain”, respectively. In particular, for mentalizing, we queried the Neurosynth database (as of Jan 2016) for published studies on the topic of “mentali*” (threshold=0.001). This resulted in a set of 112 studies with associated MNI coordinates, which we then submitted to a Neurosynth metaanalysis and saved the FDR 0.01 corrected reverse inference (RI) map. We proceeded analogously for studies of social pain (search term “social* & pain*”, 39 studies). Finally, we extracted result clusters from the “mentalizing” and “social pain” RI maps using BSPMview. In addition, we consulted two separate researcher-curated meta-analyses of “mentalizing” (60) and “social pain” (11), respectively, finding good overall agreement with the coordinates obtained via automated meta-analysis. Lastly, we discarded regions with cluster sizes below 140 mm^3^ or subcortical clusters, and symmetrized the coordinates of clearly bilateral nodes (aINS, TPJ). This procedure provided the following coordinates for the mentalizing network (see Figure S1): DMPFC (0, 53, 30), vmPFC (0, 48, −18), Precuneus (0, −54, 44), rTPJ (48, −56, 23), lTPJ (−48, - 56, 23), rMTG (53, −12, −16), and lMTG (−53, −12, −16); for the social pain network: dACC (0, 16, 32), r-aINS (38, 7, −4), l-aINS (−38, 7, −4). In addition to the regions of these two *a-priori* networks, we also examine connectivity changes during social exclusion using a whole-brain parcellation that assigns 264 brain regions to one of 13 functional networks (43).

**Figure 6.**
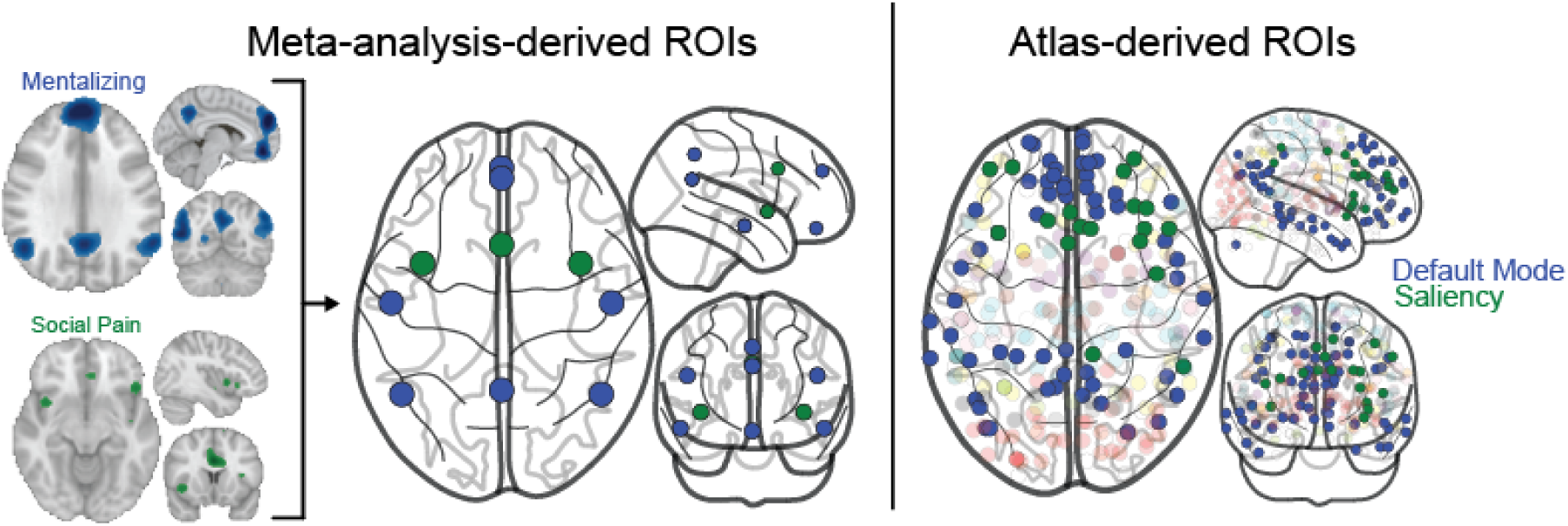
The left panel pictures the nodes for social pain and mentalizing networks as derived by synthesizing automated and researcher-curated meta-analyses on “mentalizing” (60) and “social pain” (11), respectively. The far left brain images in the panel show the meta-analytic contrast of the reverse inference maps (pFgA; FDR q= 0.01; smoothed 8 mm FWHM), which are combined to yield the parcellation shown in the schematic brain figure on the right side of the panel. The right panel pictures the regions used in the whole-brain analysis, with the default mode network colored in blue to reflect its overlap with the mentalizing network, and the saliency network colored green to capture its overlap with the social pain network.

### Functional connectivity analysis

Analysis of functional connectivity was performed in Python 2.7 using the *nilearn* package (61). Data were bandpass filtered between 0.06–0.12 Hz (21), detrended, standardized, and extracted from 8 mm radius spheres around the nodes specified above. Artifacts were reduced using frame censoring and regression. In particular, frames with framewise displacement (FD) > 0.5 mm were censored, and nuisance regressors included white matter and ventricular signals, the realignment parameters, and high-variance confounds. From the extracted time series corresponding to the social inclusion and exclusion runs, we used Pearson’s correlation coefficient to construct task-specific undirected and weighted functional connectivity matrices for each participant. To determine whether results observed could be explained by increased mind-wandering over time in the scanner, an additional set of split-half analyses were also implemented using parallel methods on the first and second half of each block (inclusion and exclusion), respectively. Analyses were implemented using python packages and in-house functions, and visualizations were created with Nilearn, Matplotlib and Seaborn (62–64).

## Acknowledgements

The authors thank Nicole Cooper, Qawi Telesford, and Marcelo Mattar for valuable comments and discussions. The authors gratefully acknowledge the University of Michigan Transportation Research Institute for research assistance; the staff of the University of Michigan fMRI Center; and Raymond Bingham, Jean Shope, Marie Claude Ouimet, Anuj Pradhan, Bruce Simons-Morton, Kristin Shumaker, Elizabeth Beard, Jennifer LaRose, Farideh Almani, and Johanna Dolle.

This research was supported by the Eunice Kennedy Shriver National Institute Of Child Health & Human Development of the National Institutes of Health under Award Number R21HD073549, a NIH New Innovator Award to E.F. (1DP2DA03515601), and by the Army Research Laboratory under Cooperative Agreement Number W911NF-10-2-0022. The views and the conclusions contained in this document are those of the authors and should not be interpreted as representing the official policies, either expressed or implied, of the NIH and its subdivisions, Army Research Laboratory, or the U.S Government. The government is authorized to reproduce and distribute reprints for governmental purposes notwithstanding any copyright notation herein. D.S.B. would also like to acknowledge support from the John D. and Catherine T. MacArthur Foundation, the Alfred P. Sloan Foundation, the Army Research Office through contract number W911NF-14-1-0679, the Army Research Office through contract number W911NF-10-2-0022, the National Institute of Mental Health (2-R01-DC-009209-11), the National Institute of Child Health and Human Development (1R01HD086888-01), the Office of Naval Research, and the National Science Foundation (CRCNS BCS-1441502, BCS-1430087, CAREER PHY-1554488.)

## Author contributions

Conceived and designed the experiments: EBF, JB, MB, CC

Performed the experiments: JB, MB, CC

Analyzed the data: RS, JB, MB

Contributed reagents/materials/analysis tools and analysis interpretation: RS, DSB, MB, JB, JG, JV, EBF

Wrote the paper: RS, EBF, JV, JG, MB, JB, DSB

## Supplementary Analyses

### Stronger Connectivity in the Mentalizing Network during Social Exclusion - Edge-level Results

The results reported in the main paper are based on the connectivity within and between the social pain and mentalizing networks, respectively. To assess results at a finer spatial resolution, we focused on individual edges and asked whether edge strength differed significantly between social exclusion and inclusion. The results of this analysis, shown in Figure S1, revealed that the edges that exhibited connectivity changes between exclusion and inclusion connected mainly regions from the mentalizing network, but not the social pain network (see Figure S1, thresholded at *p* < 0.05, uncorrected, for exploratory purposes).

**Figure S1:**
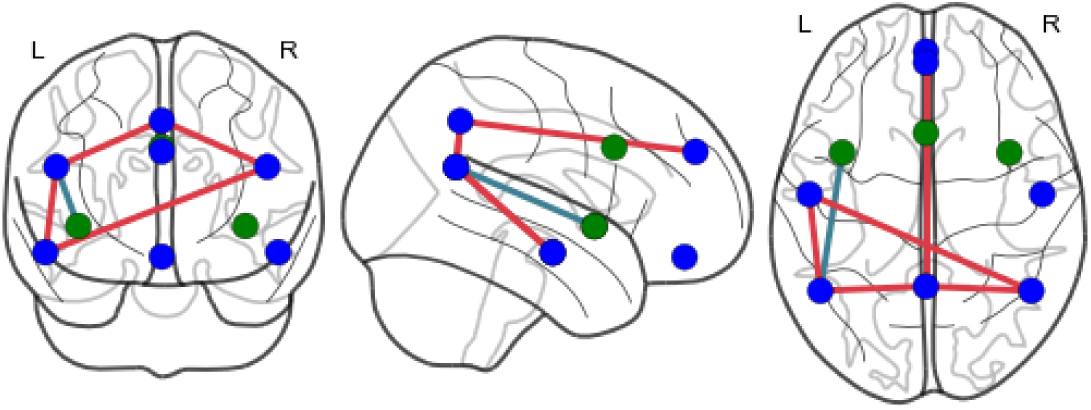
Social exclusion is associated with increased connectivity in the mentalizing network.

### Assessing the robustness of effects within the mentalizing network

A control analysis compared time courses from the first and second half of each block and confirmed increased within-network connectivity was sustained throughout the first and second halves of each block (first half: *t*(79) = 2.23; *p* = 0.029, and second half: *t*(79) = 2.57, *p* = 0.012, Figure S2)

**Figure S2:**
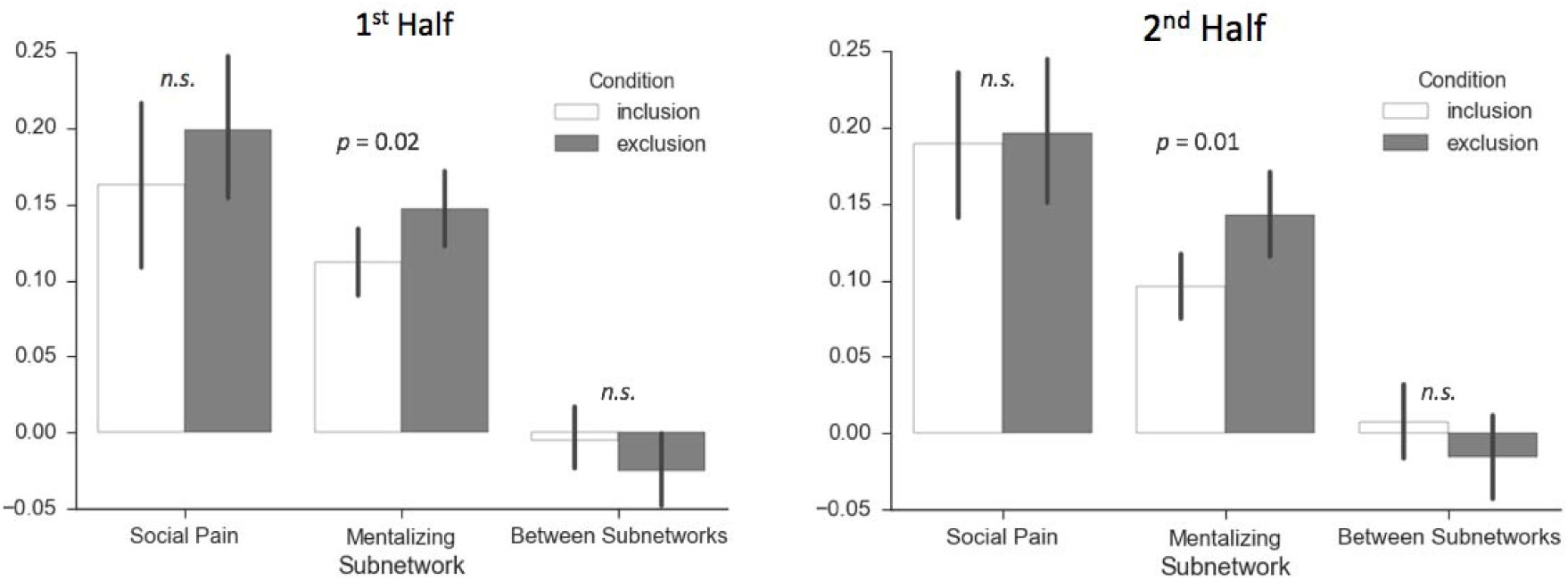
Social exclusion was associated with increased connectivity within the mentalizing network. This figure is based on the same analyses illustrated in Figure 3 in the main text, but computed using data from only the first or second half of each block, respectively.

